# Fibronectin type III and intracellular domains of Toll-like receptor 4 interactor with leucine-rich repeats (Tril) are required for developmental signaling

**DOI:** 10.1101/158782

**Authors:** Hyung-Seok Kim, Autumn McKnite, Yuanyuan Xie, Jan L. Christian

**Author notes:** To whom correspondence should be addressed: Prof. Jan L. Christian, Department of Neurobiology and Anatomy, University of Utah School of Medicine, Bldg 570 BPRB, Rm 320, 20 South 2030 East, Salt Lake City, Utah 84112.

## Abstract

Toll-like receptor 4 interactor with leucine-rich repeats (Tril) is a transmembrane protein that functions as a coreceptor for Toll-like receptors (Tlrs) to mediate innate immune responses in the adult brain. Tril also triggers degradation of the Bmp inhibitor, Smad7, during early embryonic development to allow for normal blood formation. Tril most likely plays additional, yet to be discovered, roles during embryogenesis. In the current studies, we performed a structure-function analysis, which indicated that the extracellular domain, including the fibronectin type III (FN) domain, and the intracellular domain of Tril are required to trigger Smad7 degradation in the early *Xenopus* embryo. Furthermore, we found that a Tril deletion mutant lacking the FN domain (TrilΔFN) can dominantly inhibit signaling by endogenous Tril when overexpressed in vivo. This finding raises the intriguing possibility that the FN domain functions to bind endogenous Tril/Tlr4 ligands, perhaps including extracellular matrix molecules. We also show that Tril normally cycles between the cell surface and endosomes, and that the Tril extracellular domain is required to retain Tril at the cell surface, while the intracellular domain is required for Tril internalization in *Xenopus* ectodermal explants. Using a CHO cell aggregation assay, we further show that, unlike other transmembrane proteins that contain leucine rich repeats in the extracellular domain, Tril is not sufficient to mediate homophilic adhesion. Our findings identify TrilΔFN as a valuable tool that can be used to block the function of endogenous Tril in vivo in order to discover additional roles during embryonic development.

---

Bone morphogenetic proteins (Bmps) play a critical role in specifying ventral and posterior fates during early development in all vertebrates (1). Bmps activate transmembrane serine/threonine receptors that phosphorylate the cytoplasmic proteins Smad1, 5 and 8 (2). Phosphorylated Smads (pSmad1/5/8) then recruit the co-Smad, Smad4 and translocate to the nucleus to induce target gene expression. During gastrulation, Bmps induce expression of the transcription factors *scl, cdx1* and *cdx4,* and these factors are necessary and sufficient to specify primitive erythroid fate (3,4). *Smad7* is another target gene that is induced by pSmad1/5/8 and Smad7 is a central hub for negative regulation of activated Bmp receptors (5). Smad7 recruits E3 ubiquitin ligases to activated Bmp receptors, targeting them for proteosomal degradation. Smad7 can also function in the nucleus to block transcriptional responses downstream of pSmad1/5/8.

We have recently shown that the transmembrane protein Toll-like receptor 4 (Tlr4) interactor with leucine-rich repeats (Tril) is required to augment Bmp signaling during gastrulation in *Xenopus* embryos (6). Tril promotes degradation of Smad7 protein, thereby relieving repression of endogenous Bmp signaling. This is essential to enable the mesoderm to be specified as blood. When expression of endogenous Tril is knocked down in *Xenopus* embryos using antisense morpholinos, high levels of Smad7 protein accumulate, particularly in the nuclear compartment (6). As a consequence, pSmad1/5/8 levels are reduced and development of blood is impaired. These phenotypes can be partially rescued by knocking down expression of Smad7 in Tril morphants, demonstrating that elevated Smad7 plays a causal role in the loss of Bmp signaling transduction in Tril morphants.

The mechanism by which Tril signals to degrade Smad7 is unknown. Tril was initially characterized as a co-receptor for Tlr3 and Tlr4 and is required to mediate innate immune responses in the brain of adult mammals (7-9). Tlrs bind microbial products or nucleic acids, initiating a signaling cascade involving cytoplasmic adaptors, kinases and ubiquitin ligases that generally culminates in activation of transcriptional regulators such as NF-κB, AP-1 and interferon regulatory factors (10). Recent studies suggest that Tlr signaling is also required for normal neurogenesis, axonal growth and structural plasticity in the developing and adult brain, but nothing is known about the signaling pathways that are initiated by Tlrs to mediate these non-immune effects (11). Tlrs also play additional non-immune roles in early developmental patterning and morphogenesis in insects, but evidence that vertebrate Tlr signaling is required for early development is lacking (12). It is not known whether Tril functions in concert with Tlrs in all contexts but, to date, there is no evidence that activation of Tlr signaling can promote degradation of Smad7.

In the current studies, we performed a structure-function analysis to identify domains of Tril that are required to trigger degradation of Smad7 in the context of the early embryo, and that are required for normal subcellular localization.

## RESULTS

### In vivo activity of wild type and deletion mutant forms of Xenopus Tril

We analyzed the subcellular localization and steady state levels of Smad7 in *Xenopus* embryos in which Tril expression was reduced by injection of a well characterized translation blocking antisense morpholino (MO) (6). Since we are unable to detect endogenous Smad7 using available antibodies we instead analyzed ectopically expressed epitope tagged Smad7. Two-cell embryos were injected with Smad7myc RNA (100 pg) together with control or Tril MOs (35 ng). At the midgastrula stage (st. 11), ectoderm was explanted from 10 embryos in each group for immunostaining, and an additional 15 embryos in each group were harvested for immunoblot analysis. Steady state levels of Smad7myc protein were 7-10 fold higher in Tril morphants than in controls in three independent experiments (Fig. 1A). Furthermore, Smad7myc protein accumulated predominantly in nuclei of Tril morphant embryos whereas it was diffusely localized throughout the cell and at the membrane in control embryos (Fig. 1B). These results replicate our published studies showing that endogenous Tril triggers degradation of Smad7 protein, particularly within the nuclear compartment (6).

**Figure 1.**
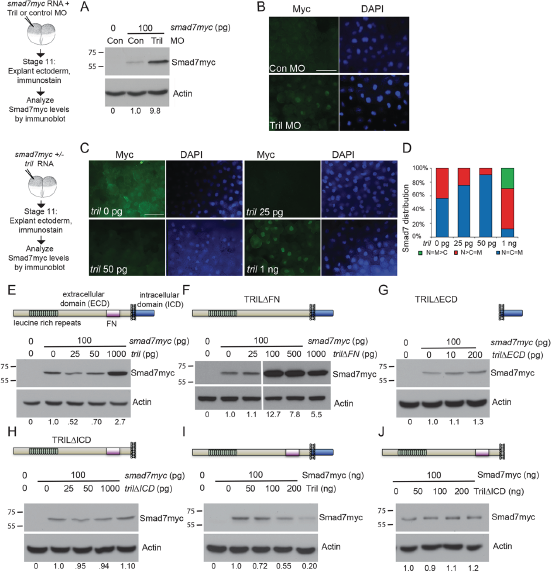
Structure function analysis of Tril. *(A,B)* Embryos were injected with RNA (100 pg) encoding Smad7myc together with control or Tril MOs (35 ng). *(A)* Immunoblots of lysates from stage 11 embryos (10 per group) were probed with anti-Myc antibodies, and then reprobed for β-Actin. Relative level of Smad7myc, normalized to actin and reported relative to that in control embryos is indicated below each lane. *(B)* Ectoderm was explanted from 5-10 embryos in each group at stage 11 and immunostained for Myc. (all images taken under identical conditions) (*C, D*) RNAs encoding Smad7-Myc alone or together with increasing doses of Tril were injected as illustrated. Ectoderm was explanted from 5-10 embryos in each group at stage 11 and immunostained for Myc. Representative immunostaining is shown in A (all images taken under identical conditions) and results are quantified in B. Results are pooled from three independent experiments. *(E-H)* RNA (100 pg) encoding Smad7-Myc was injected into two-cell embryos alone or together with RNA encoding wild type or deletion mutant forms of Tril. Immunoblots of lysates from stage 11 embryos (10 per group) were probed with anti-Myc antibodies, and then reprobed for β-Actin. The relative level of Smad7myc, normalized to actin and reported relative to that in embryos injected with Smad7myc alone is indicated below each lane. Results were replicated in three to five experiments. In panel B, all lanes are from the same immunoblot, aligned following removal of an intervening lane (following the third lane, marked by a white bar). *(I-J)* HeLa cells were transfected with DNA encoding Smad7myc alone or together with DNA encoding Tril or Tril∆ICD. Cell lysates were harvested 24 hours later and immunoblots were probed with anti-Myc antibodies, and then reprobed for β-Actin. Results were replicated in a minimum of three experiments. The relative level of Smad7myc, normalized to actin and reported relative to that in cells transfected with Smad7myc alone is indicated below each lane.

We next used a gain of function assay to identify functional domains of Tril that are required to trigger degradation of Smad7. First, we injected RNA encoding Smad7myc (100 pg) alone, or together with increasing doses of RNA encoding wild type Tril (25 pg-1 ng), into one cell of two cell embryos. At the midgastrula stage (st. 11), ectoderm was explanted from 10 embryos in each group for immunostaining, and an additional 15 embryos in each group were harvested for immunoblot analysis. In explants isolated from embryos injected with *smad7myc* RNA alone, Smad7myc was evenly distributed throughout the cytoplasm, nucleus and membrane, or was slightly enriched in the nucleus (Fig. 1C, D). In explants isolated from embryos co-injected with 25 or 50 pg of *tril* RNA, Smad7myc staining was barely detectable when imaged under identical conditions (Fig. 1C), and was evenly distributed throughout the nucleus and cytoplasm in most explants (Fig. 1D). By contrast, Smad7myc was detected primarily in nuclei in explants from embryos co-injected with 1 ng of *tril* RNA (Fig. 1C, D), similar to what is observed when expression of Tril is knocked down using antisense MOs (Fig. 1B). Immunoblot analysis revealed that steady state levels of Smad7myc were reduced in embryos co-injected with low doses of *tril* RNA (25 or 50 pg). Thus, ectopic Tril can activate signaling when expressed at low doses, triggering degradation of Smad7myc. By contrast, steady state levels of Smad7myc were enhanced in embryos injected with the highest dose of *tril* RNA (1 ng) (Fig 1E), identical to what is seen in Tril morphants (Fig. 1A and 6). This result suggests that supraphysiological levels of ectopic Tril can dominantly interfere with the ability of endogenous Tril to trigger degradation of Smad7. While this appears counterintuitive, it is possible that wild type Tril squelches signaling by disrupting the normal stoichiometry of upstream or downstream binding partners that are required to transduce signals, as has been observed for other wild type proteins (13).

We next analyzed steady state levels of Smad7myc in embryos expressing deletion mutant forms of Tril lacking putative functional domains. Tril is a single pass transmembrane protein with a large extracellular domain (ECD) that contains multiple leucine rich repeats and a fibronectin type III (FN) domain as well as a small intracellular domain (ICD) that lacks any obvious signaling motifs (illustrated above Fig. 1E). Steady state levels of Smad7myc were not reproducibly altered in embryos injected with 25 pg (Fig. 1F) or 50 pg (data not shown) of RNA encoding a Tril deletion mutant lacking the FN domain (Tril∆FN). By contrast, embryos injected with 100 pg or more of *tril ∆fa* RNA accumulated high levels of Smad7myc (Fig. 1F) in the nuclei of cells (Fig. S2), phenocopying what is seen in Tril morphants (6). Thus, the FN domain is essential for Tril function, and ectopic Tril∆FN dominantly suppresses signaling by endogenous Tril, possibly by serving as a sink for ligand or signal transduction components. We next analyzed Smad7 levels in embryos ectopically expressing mutant forms of Tril lacking the entire ECD (Tril∆ECD) or ICD (Tril∆ICD). Levels of Smad7myc were not reproducibly changed in embryos injected with 5, 10, or 200 pg of *tril∆ECD* RNA (the molar equivalent of 25, 50 or 1000 pg of wild type Tril) (Fig. 1G and data not shown), nor were they reproducibly changed in embryos injected with 25, 50 or 1000 pg of RNA encoding Tril∆ICD (Fig. 1H). Thus, the ECD and the ICD of Tril are essential for signal transduction.

Notably, injection of increasing doses of RNA encoding wild type or deletion mutant forms of epitope (HA)-tagged Tril led to a dose dependent increase in Tril-HA proteins (Fig. S1).

To ask whether the ability of Tril to destabilize Smad7 is conserved in mammals, we transfected increasing amounts of wild type Tril, or Tril∆ICD into HeLa cells. Ectopic Tril (Fig. 1I), but not Tril∆ICD (Fig. 1J), led to lower steady state levels of Smad7myc protein, but not RNA (not shown) in HeLa cells demonstrating that ectopic Tril can stimulate degradation of Smad7 in mammalian cells.

Our previous studies have shown that endogenous Tril is required for blood formation (6). As a further test of whether ectopically expressed wild type or deletion mutant forms of Tril can function as dominant mutants, we asked whether any of these proteins can phenocopy the loss of blood that is observed in Tril morphant embryos. We injected 1 ng of RNA encoding Tril, Tril∆FN or Tril∆ICD, or the molar equivalent (200 pg) of RNA encoding Tril∆ECD into each ventral blastomere near the marginal zone of 4cell embryos. We then analyzed expression of the RBC differentiation marker, *hba3.L (α-globin),* at stage 34 using either whole mount in situ hybridization (Fig. 2A, B) or qPCR analysis (Fig. 2C). Embryos injected with a total of 2 ng of *tril* or *tril∆FN* RNA showed a reproducible reduction in expression of *hba3.L* in the posterior ventral blood island (pVBI), which is the region derived from the ventral cells that were injected. By contrast, embryos injected with an equivalent dose of *Tril∆ECD* (400 pg) or *Tril∆ICD* (2 ng) RNA did not show a reproducible loss of blood (Fig. 2B-C). Notably, in 2 out of 6 experiments, embryos made to overexpress Tril∆ICD showed a partial reduction in levels of *hba3.L* relative to embryos injected with RFP, suggesting that this construct might have a mild dominant negative effect, but this was not reproduced in 4 additional experiments. Collectively, our results showing that wild type Tril, but not Tril lacking the FN domain of the ECD, or the ICD can induce degradation of Smad7 demonstrates that these domains are essential to transduce Tril signals. Furthermore, our finding that ectopic expression of high doses of wild type Tril, or Tril∆FN phenocopies the accumulation of Smad7myc and loss of blood that is observed in Tril morphants demonstrates that these constructs can dominantly block endogenous Tril function.

**Figure 2.**
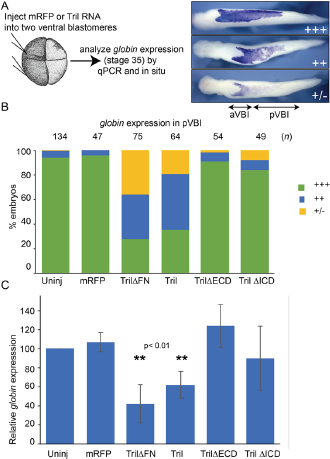
Overexpression of Tril or Tril∆FN inhibits blood formation. *(A,B)* RNA encoding mRFP (1.5 ng) or wild type or deletion mutant forms of Tril was injected into both ventral cells of 4-cell embryos and expression of *hba3.L* was analyzed by WMISH at stage 34 in three experiments. hba3.L staining in the posterior VBI (pVBI) was scored as absent or very weak (+/-), weak (++) or strong (+++), as illustrated. aVBI, anterior VBI. Results are pooled from a minimum of three independent experiments. *(C)* RNA encoding mRFP (1.5 ng) or wild type or deletion mutant forms of Tril was injected into both ventral cells of 4-cell embryos and expression of *hba3.L* was analyzed by qPCR at stage 34. Mean ± s.d. are shown, n=4, ** indicates p < 0.01 by two-tailed t-test.

### Tril localizes to the plasma membrane and intracellular vesicles in Xenopus embryos, but is found mainly in endosomes in HeLa cells

We next examined the subcellular localization of Tril in *Xenopus* embryos. We and others (8) have been unable to generate antibodies that recognize endogenous Tril and thus we injected RNAs encoding HA epitope tagged Tril (100 pg) together with membrane tagged RFP (memRFP) (50 pg) into a single blastomere at the four-cell stage. Ectoderm was explanted from embryos at midgastrula stage 11 and immunostained using antibodies specific for HA and RFP (illustrated in Fig. 3A). Tril was present primarily at the plasma membrane in *Xenopus* explants but was also readily detected in intracellular vesicles (Fig. 3B). By contrast, in transiently transfected HeLa cells, ectopic Tril HA was weakly detected at the cell surface but was more abundant in intracellular vesicles that partially co-localize with the early endosomal marker Rab5 (Fig. 3C).

**Figure 3.**
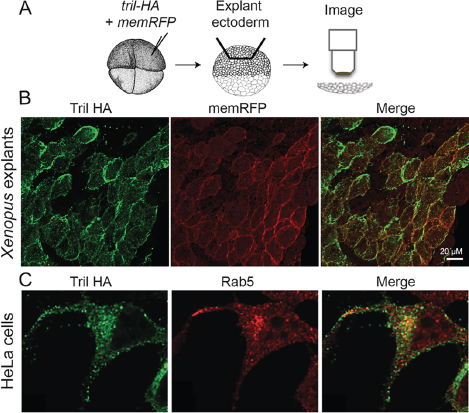
Tril localizes to the plasma membrane and intracellular vesicles in *Xenopus* embryos, but is found mainly in endosomes in HeLa cells. *(A-B)* RNAs encoding Tri-HA and membrane-targeted RFP (memRFP) were injected into a single animal pole blastomere of 4-cell embryos as illustrated. Ectoderm was explanted from 5-10 embryos in each group at stage 11 and immunostained for HA and RFP. *(C)* HeLa cells were transfected with DNA encoding Tril-HA. Cell were harvested 24 hours later and double label immunostained with antibodies specific for HA and Rab. Results were replicated in three experiments.

We used an antibody uptake assay (14) to test whether the Tril protein detected in intracellular vesicles is retrieved from the cell surface as opposed to being present in vesicles enroute to the plasma membrane. HeLa cells were transiently transfected with DNA encoding Tril containing an HA epitope tag in the ECD (illustrated above Fig. 4, yellow bar) and a Flag epitope tag at the end of the ICD (red bar). After 24 hours, cells were permeabilized and double label immunofluorescence was used to detect the HA and Flag epitopes. Under steady state conditions, both antibodies recognize ectopic Tril near the cell surface and in vesicles throughout the cell (Fig. 4). For the uptake assay, cells were incubated at 4° C for one hour together with antibodies specific for the HA tag. This allows the HA antibody to bind to the extracellular domain of Tril molecules that are located on the cell surface. Cells were then washed to remove unbound antibody and were fixed immediately (0 minutes), or were incubated at either 4° C (which prohibits endocytosis) or at 37° C (which is permissive for endocytosis) for one hour before fixation. Cells were then permeabilized, incubated with antibodies specific for the Flag tag, and the Flag and HA epitopes were visualized using Alexa-488 and Alex-568 coupled secondary antibodies (illustrated above Fig. 4). Immediately after incubation at 4° C (Fig. 4B-B’’) or following an additional one hour incubation 4° C (Fig. 4C-C”) the HA epitope tag was detected only at the cell surface, demonstrating that the block to endocytosis was complete, while the Flag epitope tag was detected primarily intracellularly. After 60 minutes at 37° C, the HA antibody bound to Tril was no longer detected at the cell surface but was instead present in vesicles that were also detected using the Flag antibody (4D-D’’). Thus, Tril is trafficked to the plasma membrane of cells and is retrieved into endosomal compartments by endocytosis.

**Figure 4.**
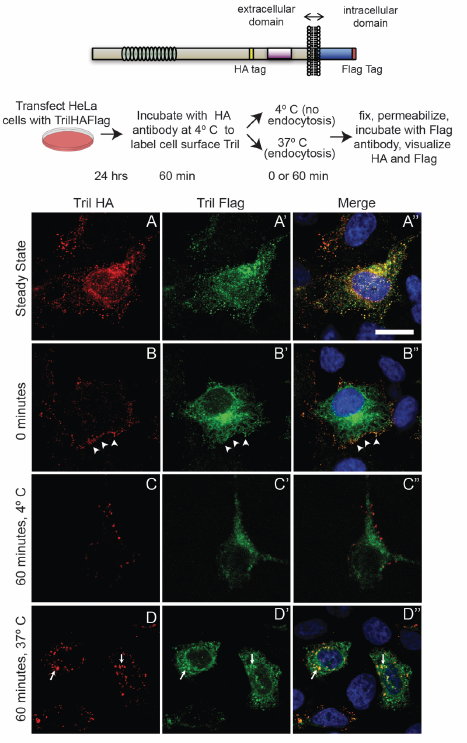
Tril is retrieved from the cell surface into endosomes in HeLa cells. Hela cells were transiently transfected with DNA encoding HA-and Flag-epitope tagged Tril (illustrated above figure) and were either permeabilized and double label immunostained with antibodies specific for both epitopes *(A-A’’)* or were incubated at 4° C with anti-HA antibodies for one hour and then either permeabilized immediately *(B-B’ ‘),* or incubated at 4° C *(C-C’’)* or at 37° C *(D-D”)* for one hour before permeabilization and addition of antibodies specific for the Flag tag. HA and Flag epitopes were visualized with species specific fluorescent secondary antibodies. The while arrowheads in B-B’ denote Trial present at the cell surface while the white arrows in D-D’ indicate Tril present in intracellular compartments. Results were replicated in three independent experiments.

### The extracellular domain of Tril is required to retain it at the plasma membrane while the intracellular domain is required for internalization

To identify domains of Tril that are required for intracellular trafficking, and to begin to ask whether Tril activity correlates with membrane and/or endosomal location in the context of the early embryo, we examined the subcellular localization of HA-tagged wild type and deletion mutant forms of Tril in *Xenopus* ectodermal explants. RNA encoding Tril variants (100 pg) was injected together with membrane tagged RFP (memRFP) RNA (50 pg) into a single animal pole blastomere of four-cell stage embryos. Ectoderm was explanted at stage 11 and immunostained with antibodies specific for the HA epitope and RFP (illustrated in Fig. 5A). As described previously, wild type Tril was strongly detected at the plasma membrane, but was also present in cytoplasmic puncta (Fig. 5B-B’’, arrowheads). By contrast, Tril∆ICD was detected only at the plasma membrane (Fig. 5C-C’’), suggesting that the ICD is required for internalization of wild type Tril. Surprisingly, Tril∆ECD was detected almost exclusively in intracellular puncta and not at the cell surface (Fig. 5D-D’’) despite the presence of an intact transmembrane domain. Tril∆FN was abundantly detected in cytoplasmic puncta, diffusely throughout the intracellular space and also at the plasma membrane, although membrane staining appeared less intense than that of wild type Tril (Fig. 5E-E’’). These results suggest that the ICD of Tril is required for internalization of the wild type protein and that the ECD is required to stably retain Tril at the cell surface.

**Figure 5.**
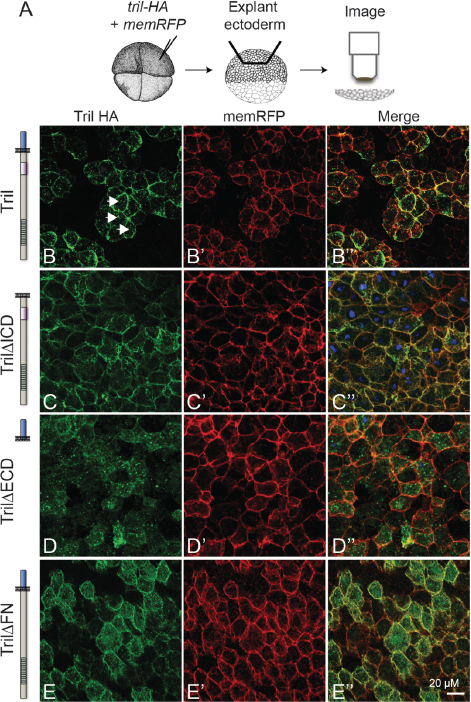
The extracellular domain of Tril is required to retain it at the plasma membrane while the intracellular domain is required for internalization. *(A-E)* RNAs encoding wild type or deletion mutant forms of Tri-HA (illustrated to left of each panel) and membrane-targeted RFP (memRFP) were injected into a single animal pole blastomere of 4-cell embryos as illustrated. Ectoderm was explanted from 5-10 embryos in each group at stage 11 and immunostained for HA and RFP. Arrowheads in A indicate intracellular staining. Results were replicated in three independent experiments.

*The extracellular domain of Tril does not promote homophilic or heterophilic aggregation in CHO cells.* The observation that the ECD of Tril is required to retain it at the cell surface raises the possibility that it does so through homophilic interactions with the ECD of other Tril molecules, or through heterophilic interactions with other cell surface proteins. Consistent with this possibility, the ECD of several other LRR-containing proteins, including Tlrs, have been shown to interact, and to mediate homophilic or heterophilic adhesion (15-18). To test whether Tril can mediate cell adhesion, we transiently transfected nonadherent CHO cells with Tril-GFP, Cadherin6-GFP (as a positive control) or GFP (as a negative control) and compared the ability of cells to aggregate. As shown in Fig. 6, only cells expressing Cadherin6 formed aggregates. Thus, the ECD of Tril does not promote homophilic adhesion, nor does it promote heterophilic adhesion with endogenous proteins expressed on the surface of CHO cells.

**Figure 6.**
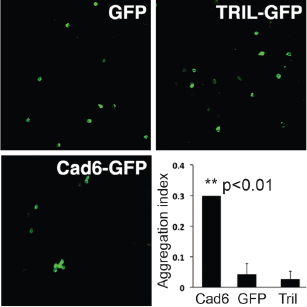
Tril is not sufficient to mediate homophilic adhesion. CHO cells transfected with either GFP (negative control), Cadherin6-GFP (Cad6-GFP) (positive control) or Tril-GFP were tested for adhesion. Only cells expressing Cadherin6 formed aggregates. The aggregation index was calculated by dividing the total GFP fluorescence in cell aggregates by the total GFP fluorescence in the well. Mean ± s.d. are shown, n = 3, ** indicates p < 0.01 by two-tailed t-test.

## DISCUSSION

In the current studies, we identified domains of Tril that are required to trigger degradation of Smad7 and thereby augment Bmp signaling to allow for normal blood formation in early embryos, and then correlated these domains with those required to for normal subcellular localization. We find that the ECD and the ICD of Tril are both required to transduce signals necessary for targeted degradation of Smad7. Furthermore, we show that the ECD is required for cell surface retention while the ICD is required for internalization.

In innate immunity, Tril functions as an accessory molecule to enhance signaling downstream of Tlr3 and Tlr4 (7,8). A variety of accessory molecules have been identified that are required to enable Tlrs to signal, and these function to deliver and enhance the affinity of ligands to their cognate receptors, but they can also act as chaperones, to traffic receptors to signaling competent subcellular locations (19). Tril was originally identified as an adaptor protein required for Tlr4 signaling. In vitro studies showed that the extracellular domain of Tril interacts directly with Tlr4, and with its ligand, LPS, suggesting that it functions primarily to enhance ligand/receptor affinity (7). Subsequent in vitro studies showed that Tril plays a similar role in regulating Tlr3 signaling, and in vivo studies confirmed that Tril is essential for maximal Tlr3-and Tlr4-dependent cytokine production in the brain in response to bacterial infection (8,9).

It is unknown, at present, whether Tril functions independently or whether it serves as a co-receptor for Tlrs in the context of early development, as it does in the immune system. While Tlrs are known to play non-immune roles in developmental patterning and morphogenesis in insects, evidence that vertebrate Tlr signaling is required for early development is lacking (12). Recent studies, however, have shown that Tlr signaling is required for normal neurogenesis, axonal growth and structural plasticity in the developing and adult brain (11), where Tril is highly expressed (9). Very little is known about the signaling pathways that are initiated by Tlrs to mediate these non-immune effects.

In the context of innate immunity, Tril appears to be required for the initial step of ligand binding and Tlr activation, but subsequent signal transduction from the membrane to the nucleus may occur independent of Tril. By contrast, our structure/function analysis shows that the ICD of Tril is indispensable for downstream signaling leading to degradation of Smad7 both in *Xenopus* and in cultured mammalian cells. Furthermore, we show that the ICD is also required for internalization of Tril. The ICD of Tril is highly conserved across species; *Xenopus* and human Tril share 77% amino acid identity in this domain. Although the ICD of Tril lacks any recognizable signaling motifs, it does include one, and possibly two conserved di-leucine like motifs ([D/E]XXL[L/I]) that may function to direct clathrin-mediated endocytosis (20). Our finding that the ICD of Tril is required for signal transduction raise the possibility that the ICD binds to adaptor proteins that directly transmit signals required for degradation of Smad7, independent of Tlrs. Alternatively, it is possible that the ICD functions only to mediate endocytic retrieval, and that Tril can only transduce signals from internal compartments within the cell. The adaptor protein CD14 controls the LPS-induced endocytosis of Tlr4 (21), which is critical to activate signaling networks culminating in production of type I interferons (22). It is possible that Tril plays an analogous role in mediating internalization of Tlrs, and that this is required to direct degradation of Smad7 during development.

Although the ICD of Tril is required to trigger degradation of Smad7, it is not sufficient to initiate signaling since Tril∆ECD, which has an intact ICD but is not stably retained at the plasma membrane, is unable to signal. It is possible that internalization serves to target the protein for degradation to terminate signaling, and that Tril∆ECD is inactive because it is rapidly degraded. This possibility seems less likely, however, because steady state levels of Tril∆ECD are equivalent to or higher than those of wild type Tril in HeLa cells or in embryos expressing equivalent amounts of each DNA or RNA, despite the fact that it is not retained at the cell surface. An alternate explanation for the lack of activity of Tril∆ECD is that signaling downstream of Tril requires receptor activation by cell surface or extracellular ligands that bind to the ECD.

Our finding that Tril∆FN is unable to initiate signaling, but can dominantly inhibit signaling by endogenous Tril suggests a critical role for the fibronectin type III domain in receptor activation. Fibronectin type III repeats are immunoglobulin like-domains that were originally identified in fibronectin, but homologous motifs have been found in a wide variety of cell surface, extracellular matrix (ECM) and cytosolic proteins (23). Functions of fibronectin type III domains are not well understood but can include interactions with other ECM or cell surface proteins (24). Interestingly, there is growing evidence that Tlrs respond to a variety of endogenous ligands. Tlr4, in particular, can be activated by ECM degradation products including fibronectin and hyaluronin (25), both of which are present in the early embryo. It is possible that the FN domain of Tril binds to ECM components that function as Tlr4 ligands, and that Tril functions as a bridge to enhance ligand/receptor affinity in the embryo as it does in the innate immune system. This scenario provides a possible explanation for the mechanism by which Tril∆FN dominantly inhibits signaling by endogenous Tril. If the ectodomain of Tril∆FN can interact with Tlr4 but cannot interact productively with ligands required to initiate signaling, this could potentially sequester endogenous Tlrs at the membrane in an inactive conformation. Alternatively, the intact ICD of Tril∆FN might sequester cytoplasmic effectors at the membrane and/or in inappropriate subcellular compartments so that they are not available to transduce signals downstream of endogenous Tril.

The mechanism by which wild type Tril dominantly inhibits signaling by endogenous Tril is more difficult to model. The most likely explanation is that signal transduction requires a specific stoichiometry of extracellular and intracellular binding partners, and that an excess of one component creates a stoichiometric imbalance within the protein complex such that signal transduction cannot proceed. Many examples of abnormal phenotypes caused by overexpression of wild type gene products involved in macromolecular complexes exist (13). The observation that neither Tril∆ICD nor Tril∆ECD can dominantly inhibit signaling by endogenous Tril is consistent with these models of squelching of intracellular or extracellular signaling components.

Previously published studies, along with our current findings, reveal that Tril is present primarily at the plasma membrane in stably transfected HEK cells (8) and in intact *Xenopus* ectodermal explants, whereas it is present primarily in endosomes in HeLa cells, or in U373 cells (8). This raises the question of what mediates cell surface retention versus internalization in these different cell types. The observation that Tril∆ECD is largely absent from the cell surface in *Xenopus* ectodermal explants, along with our data showing that Tril does not mediate homophilic adhesion, suggests that interactions between the ECD of Tril and heterologous cell surface proteins and/or components of the ECM are required to prevent constitutive internalization in the absence of pathway stimulation. Future studies are aimed at identifying the proteins that interact with, and transduce signals downstream of Tril during early development.

## EXPERIMENTAL PROCEDURES

*Xenopus embryo culture and manipulation*-Animal procedures followed protocols approved by the University of Utah Institutional Animal Care and Use Committee. Embryos were obtained, microinjected, and cultured as described (26). Embryos were processed for in situ hybridization as described (27) except that the vitelline coat was not removed prior to fixation and BM purple (Roche) was used as a substrate.

*cDNA constructs and* MOs-Sequence encoding an HA epitope tag was inserted into the ECD of *Xenopus tril.S* between amino acids 499/500 to generate Tril-HA and/or sequence encoding a Flag epitope tag was appended following the last amino acid to generate Tril-HAFlag/Tril-Flag using a PCR based approach. Tril-HAFlag can rescue blood formation in embryos in which translation of endogenous Tril is blocked, demonstrating that the epitope tags do not interfere with Tril function (6). cDNAs encoding deletion mutant forms of *tril* were made using the splicing by overlap extension method (28) or by PCR-mediated introduction of appropriate restriction sites. Tril∆FN lacks amino acids 650-730, Tril∆ECD lacks amino acids 45744 and TrilACT lacks amino acids 792-863 of Tril. Previously characterized Tril MOs (6) were purchased from Gene Tools, LLC.

*Immunoblots-Embryos* were lysed in Triton X-100 lysis buffer (29) containing HALT protease inhibitor and phosphatase cocktail (Thermo Scientific). Immunoblots were performed as described (30) with the following antibodies: Rat anti-HA (Roche Rat monoclonal 3F10, cat. No. 11 867 423 001, lot # 12213, 1:1000), Mouse anti-Myc (Developmental Hybridoma bank, 9E10, 1:1,000) and Rabbit anti-p-Actin (Abcam ab82272, lot # GR265013-1, 1:10,000). Uninjected embryos were included on all anti-myc blots as a negative control to ensure antibody specificity.

*Immunostaining*-Ectodermal explants were immunostained with antibodies specific for HA (Roche Rat monoclonal 3F10, cat. No. 11 867 423 001, lot # 12213, 1:500) or myc (Cell Signaling, 7D10 Rabbit mAb #2278, lot # 5, 1:250) as described (6). Hela cells were transiently transfected with 10 ng of DNA encoding wild type or deletion mutant Tril-HAFlag in pCS2+. After 24 hours, cells were fixed in 4% paraformaldehyde/PBS for 5 minutes and then immunostained with antibodies specific for Flag (Sigma mouse mAb clone M2, F1804, lot # SLBK1346V, 1:500) and/or HA (Roche, Rat monoclonal 3F10, cat. No. 11 867 423 001, lot # 12213, 1:500). Primary antibodies were detected with alexa 488-conjugated anti-rabbit (Invitrogen, A11008, lot # 1408830) or anti-rat (Invitrogen, A21208, lot #1701951), or alexa 568-conjugated anti-mouse (Invitrogen, A11031, lot # 6592-1) secondary antibodies used at a dilution of 1:1000.

For antibody uptake assays, transfected cells were incubated with 1:200 Rat anti-HA antibody for one hour at 4° C and then were either fixed prior to addition of anti-Flag antibodies, or were incubated for an additional 60 minutes at 4° or 37° C prior to fixation and immunostaining with Flag as described above. To detect Smad7myc, explants were imaged using a Leica DM2500 compound microscope and a Leica DFC425 C digital camera. For all other immunostaining, explants and cells were imaged using an Olympus FV1000 confocal microscope.

*Analysis of RNA*-Total RNA was isolated and qPCR was performed as described (31) using an annealing temperature of 58°C. Primer sequences have been reported previously (6).

*CHO cell aggregation assay*-CHO cells were transfected with hTril-GFP/pCDNA3.1 (5 μg), Cadherin6-GFP/pCAG (5 μg), or GFP/pCS2+ (5 μg) using PEI (polyethylenimine). 48 hrs. later, cells were washed with HEPES Mg^2+^ free (HMF) buffer (137 mM NaCl, 5.4 mM KCl, 1 mM CaCl2, 0.34 mM Na2HPO4, 5.5 mM glucose, 10 mM HEPES, pH 7.4 adjusted with NaOH) and detached from the dishes using 0. 01% trypsin in HMF. Detached cells were spun down, resuspended in HMF, counted, and 100,000 cells were pipetted into single wells of 24-well plates precoated with 1% BSA in HMF. Subsequently, the plates were placed on a nutator for 90 min at 37°C. The cells were then fixed with paraformaldehyde (PFA) (4% final concentration), transferred to a 96-well glass bottom plate, and imaged using an Olympus FV1000 confocal microscope. The aggregation index was calculated by dividing the total GFP fluorescence in cell aggregates by the total GFP fluorescence in the well. Analysis was done using ImageJ.

***Statistical analysis***-NIH Image J software was used to quantify band intensities. A student’s *t*-test was used to compare differences between two groups. Differences in subcellular localization of Smad7 were analyzed using GraphPad software to conduct two-way ANOVA followed by a Bonferroni multiple comparisons test. Differences with P < 0.05 were considered statistically significant.

## Acknowledgements

We thank Anne Martin for help with troubleshooting the adhesion assay in CHO cells and Dr. Luke O’Neill for the human Tril-GFP construct.

## Author Contributions

HK conducted Western analysis, performed confocal microscopy of explants and is solely responsible for the results shown in Fig. 4 and Fig. 6, AM conducted Western analysis, in situ hybridizations and qPCR analysis of *globin* expression, YX conducted exploratory immunostaining and embryo injection experiments and provided the results shown in Fig. 3C, JC conceived the idea for the project, performed all embryo injections, generated and immunostained embryonic explants and wrote the paper.

## FOOTNOTES

This work was supported by the National Institute of Child Health and Human Development [R01HD067473 to J.L.C], the Huntsman Cancer Foundation [150203 to JLC] and the National Cancer Institute [P30CA042014]. This work utilized DNA, peptide and imaging shared resources supported by the Huntsman Cancer Foundation and the National Cancer Institute [P30CA042014]. The content is solely the responsibility of the authors and does not represent the official views of the National Institutes of Health.

